# Shear stress mediates metabolism and growth in electroactive biofilms

**DOI:** 10.1101/391029

**Authors:** A-Andrew D Jones, Cullen R Buie

## Abstract

Electroactive bacteria such as *Geobacter sulfurreducens* and *Shewanella onedensis* produce electrical current during their respiration; this has been exploited in bioelectrochemical systems. These bacteria form thicker biofilms and stay more active than soluble-respiring bacteria biofilms because their electron acceptor is always accessible. In bioelectrochemical systems such as microbial fuel cells, corrosion-resistant metals uptake current from the bacteria, producing power. While beneficial for engineering applications, collecting current using corrosion resistant metals induces pH stress in the biofilm, unlike the naturally occurring process where a reduced metal combines with protons released during respiration. To reduce pH stress, some bioelectrochemical systems use forced convection to enhance mass transport of both nutrients and byproducts; however, biofilms’ small pore size limits convective transport, thus, reducing pH stress in these systems remains a challenge. Understanding how convection is necessary but not sufficient for maintaining biofilm health requires decoupling mass transport from momentum transport (i.e. fluidic shear stress). In this study we use a rotating disc electrode to emulate a practical bioelectrochemical system, while decoupling mass transport from shear stress. This is the first study to isolate the metabolic and structural changes in electroactive biofilms due to shear stress. We find that increased shear stress reduces biofilm development time while increasing its metabolic rate. Furthermore, we find biofilm health is negatively affected by higher metabolic rates over long-term growth due to the biofilm’s memory of the fluid flow conditions during the initial biofilm development phases. These results not only provide guidelines for improving performance of bioelectrochemical systems, but also reveal features of biofilm behavior. Results of this study suggest that optimized reactors may initiate operation at high shear to decrease development time before decreasing shear for steady-state operation. Furthermore, this biofilm memory discovered will help explain the presence of channels within biofilms observed in other studies.

---

Bacteria exist in biofilms more than in planktonic states to (1) protect bacteria from predation and chemical attack and (2) to allow bacteria to manipulate their environment.^1^ Electrochemically active biofilms^2-4^ not only have potential to bridge the energy-water nexus, producing power from wastewater treatment,^5,6^ but also to be used as a tool to study biofilm development because they produce a measurable current as part of their metabolism.^2^ Though bioelectrochemical systems employ flow to improve nutrient delivery and metabolic waste removal,^6-9^ which is known to influence biofilm behavior, the influence of shear on electrochemically active biofilms remains largely unexplored.^6-10^ That fluid shear affects bacteria biofilms fixed to a surface and that biofilms have channels^11^ is not well understood because pore sizes limit convection^12-15^ and the influence of shear. Electroactive bacteria form thicker biofilms^5^ and a larger portion stay metabolically active^16,17^ than soluble respiring bacteria and should therefore be more susceptible to fluid shear. Using the analytic equations for flow and mass flux that exist for a rotating disk electrode, the absolute concentration of nutrients is reduced in a way that decouples the influence of mass and momentum flux. In doing so, we find operating conditions that may improve the performance of bioelectrochemical systems. In addition, we demonstrate that while these biofilms are affected by their environment, their porosity does not change at steady-state so they are still susceptible to decay from pH stress.^5^

We used a rotating disk electrode that has analytic equations for flow and mass flux that can be varied independently, and through scaling laws, have been used to simulate pipe flow, conduit flow and spray jets.^18^ The property of separation, specifically, makes rotating disk electrodes useful for studying electrochemical reactions; however, when it has been used to study electroactive biofilms in the past, this particular property was not used.^19^ We benchmarked our initial flux and shear to that of a practical bioelectrochemical system,^6^ and varied the shear by two orders of magnitude. We used a pure culture of *Geobacter sulfurreducens*, as it has been shown to produce the highest current^5^ in a bioelectrochemical system and is the most abundant species in mixed-culture bioelectrochemical systems.

## Methods

### Practical Benchmark

We benchmarked this study to an upflow microbial fuel cell by He et al.^6^ An upflow microbial fuel cell is designed in a similar manner to an upflow anaerobic sludge blanket bioreactor where organic laden fluid is flown upwards over a biofilm-coated porous media, having a smaller footprint than continuous stirred tank reactors and subsequently higher shear.^20^ Matching two fluid flow systems using only the dimensionless Reynolds number that is a ratio of momentum fluxes, produces vastly different results between the two systems.^21^ Instead, both the mass flux and the momentum flux must be matched. The dimensionless shear stress for the upflow microbial fuel cell is found using SI Eq. 5, 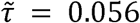 ̃ 0.056. This value is matched to the shear of rotating disc setups which for our system results in a rotation rate of 739 rpm, a shear stress of 1.048 Pa and shear rate of 1046 s^−1^. Using the data from He et al.,^6^ and SI Eq. 6 the flux can be found *N* = 5.35 e - 6 g COD cm^−2^ s^−1^ We varied the dimensional shear stress over 2 orders of magnitude using the keeping the mass flux fixed resulting in the concentrations, rotation rates, and dimensionless shear stress in Table 1.

**Table 1:** The experimental parameters used for an acetate-fed *Geobacter* rotating disk. The top row is based on the experimental parameters of He et al., scaled as described herein.^6^

| Rotation Rate [RPM] | Shear [Pa] | Concentration [mM] | Dimensionless Shear [] |
| --- | --- | --- | --- |
| 739 | 1.048 | 14.97 | 0.0562 |
| 159 | 0.1048 | 32.24 | 0.1211 |
| 34 | 0.01048 | 69.47 | 0.2610 |

The current will be normalized to the mass transfer limiting current for a rotating disk,

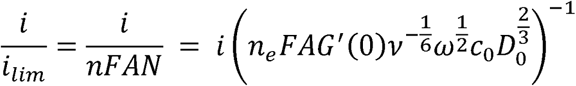

To correlate this with the flux through the biofilm, the growth time of the biofilm is taken into account as

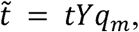

where *Y* is the biomass yield of the substrate and *q*_*m*_ is the maximum substrate utilization rate^22^. A plot of 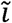 versus 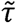 at constant mass flux results in curve that is similar to that of logistic growth^23^ with parameters that may relate to physical mechanisms.

### Media

The media was 0.59 g potassium dihydrogenphosphate, 0.38 g potassium chloride, 2.19 g sodium hydrogen carbonate, 0.36 g sodium chloride, 0.20 g ammonium chloride, 0.04 g calcium chloride dihydrate, and 0.10 magnesium chloride hexahydrate (MilliporeSigma, Darmstadt, GER) in 1 L of deionized water based on the media used in the literature. ^19,24^ The resulting conductivity was 2.10 mS.cm^−1^ at 30 ºC. To this media was added 14.97 mL, 32.24 mL, or 69.47 mL of 1 M sodium acetate in the same media. This resulted in conductivity between 5.5 to 6.7 mS.cm^−1^. Acetate concentration was held constant by removing old media and adding gas sparged new media added using a peristaltic pump at 0.07 mL. min^−1^.

### Electrodes

The working electrode was a 0.4 cm diameter graphite exchange disk (Pine Instruments, Durham, NC, USA). The electrode was soaked in 1 M hydrochloric acid and 1 M sodium hydroxide for 24 hours after wiping with ethanol and roughened using 5 μm grit silicon carbide polishing paper to clean from previous experiments. A single junction Ag/AgCl (3.8 M KCl) reference electrode (MilliporeSigma, Darmstadt, GER) was used because the conductivity of the media is too low to sustain a double-junction. While this may cause leakage of the 3.8 M sodium solution into the electrolyte, the abiotic controls do not show any significant deviation. A 0.1 cm diameter graphite electrode behind a glass frit was used as the counter electrode (Pine Instruments, Durham, NC, USA).

### Apparatus

The reactor was a water jacketed inverted 500 mL Erlenmeyer flask, with 4 ports for electrodes and gas (Pine Instruments, Durham, NC, USA). The reactor was maintained at a constant temperature of 30 ºC using a water jacket (Julabo E5, Julabo USA, Allentown, PA, USA). The system was purged and blanketed at 50 sccm with ultra-high purity 80% nitrogen and 20% carbon dioxide gas (Airgas Inc, Radnor Twp, PA, USA). The current was measured with a Gamry Reference 600 or a Gamry Reference 3000 (Gamry Instruments, Philadelphia, PA, USA) with a resolution of 600 mA and a bandwidth of 10 MHz which is much lower than the sampling rate and current measured.

### Bacteria

*Geobacter sulfurreducens* PCA (ATCC, 51573), was grown anaerobically from a frozen stock in a media matching Coppi et al. with 4 mM L-cystine and 40 mM fumarate^24^ at 30 ºC. This was transferred to a 100 mL flask and grown for 2 days. The cells were centrifuged twice at 6000 rpm for 10 min rinsed in between with the media described earlier. Optical density was measured at 600 nm (UV-1800, Shimadzu, Nakagyo-ku, Kyoto, JPN) after suspending the cells in the full 100 mL. Cell density was then calculated using a disposable hemocytometer (Incyto, Co., Ltd, Chonan-si, Chungnam-do, KOR). Cell volume was calculated assuming the bacteria were cylindrical and measuring them using a 100 X oil immersion objective.

### Procedure

The media was brought to 30 ºC and internal resistance of the media was measured. All conditions were run in biological triplicate. A roughly 24 hr cycle included 30 min open circuit potential measurement, followed by two cycles of cyclic voltammetry, chronoamperometry for 22 hours, and electrochemical impedance spectroscopy. Chronoamperometry was conducted at −0.156 V vs Ag/AgCl based on Soussan et al.^25^ Open circuit potential was measured at a sampling rate of 0.016 Hz for 30 min as this appeared sufficient to reach steady-state. Cyclic voltammetry had an equilibration time of 5 s, a step size of 2 mV and a scan rate of 1 mV.s^−1^ from −0.755 – 0.045 V. following the method of Marsili et al.^26^ The electron diffusion current was measured as the maximal current generated during a slow (2 mV.s^−1^) cyclic voltammetry scan.

Electrochemical impedance of the biofilm is measured since it may be related to the structure of the biofilm and is a key characteristic of microbial fuel cell performance. ^12,27,28^ Electrochemical impedance spectroscopy was conducted using two sets of parameters 1000000 Hz to 0.10 Hz at *E*_*dc*_ = −0.340 V, *E*_*ac*_ = 5 mV and 1000000 Hz to 0.01 Hz at *E*_*dc*_ = −0.157 V, E_ac_ = 10 mV following both Babauta and Beyanal and Marsili et al. ^19,26^ However, there is no accepted model for biofilm impedance so we do not fit the impedance spectra.

The biofilm density and thickness are necessary to compare the results to previous research on bioreactors. ^29^ After 7 days, the bacteria on disk were dyed using 4’,6-diamidino-2-phenylindole a DNA stain (NucBlue, Molecular Probes, Inc, Eugene, OR, USA) and Alexa Fluor^®^ 594 - Concanavalin A, Conjugate (Molecular Probes, Inc, Eugene, OR, USA) that binds to glycocalyx. 14 After 10 minutes, the cells were fixed for 24 hrs using 4 % SEM grade glutaraldehyde (MilliporeSigma, Darmstadt, GER) solution in their native buffer. The biofilm was then rinsed in an ethanol series 30, 50, 70, 90, 100, 100. These were imaged using confocal microscopy on four 50 μm^2^ regions at the perimeter of the disk where the shear was calculated. The sample was then critical point dried and sputter coated. The images were processed in MATLAB where a threshold was set and the images were converted to black and white. Porosity was calculated layer by layer.

## Results and Discussion

Increasing shear stress from 0.01 Pa to 1 Pa reduced the doubling time of the current from 0.26 days to 0.14 days, as shown in SI Figure S1 and Table 2. The near immediate current response is found in the literature,^30^ but here we also found a dependence on shear stress. We propose that the rapid development of measurable current is also independent of the flux of bacteria, which was not held constant. If the variable flux of bacteria caused the rapid current development, it would be reflected in the initial average open circuit potential (OCP) which is used to describe active bacteria colonization of an electrode.^31^ However, the OCP shows ~25% drop (Figure 1b) that is inconsistent with the trend in current rise. We assume that all the bacteria had the same initial electrochemical activity, as they were all taken from mid-log planktonic growth phase^17^ and seeded at 600 RPM for 10 minutes prior to the start of each experiment. However, the variability shown in Figure 1b demonstrates that further study of the influence of bacteria on electrode potential is needed.^31^

**Table 2:** Growth and current parameters show linear dependence on shear stress. There are linearly increasing trends in doubling time and time to maximum current with increased shear, and linearly decreasing trends in maximum current. The time, *t*_*max*_, to maximal growth is estimated using a dimensionless model based on logistic-growth (*R*^2^ > 0.9). The doubling time of current, *t*_1/2_ estimated assuming an exponential rise over the first 24 hrs.

| Shear [Pa] | $t_{1/2}$[days] | $t_{max}$[days] | $I_{max}$ [μA] |
| --- | --- | --- | --- |
| 1 | 0.14 | 3.8 | 136±1.24 |
| 0.1 | 0.12 | 4.3 | 89.3±1.24 |
| 0.01 | 0.26 | 9.8 | 54.3±7.08 |

**Figure 1.**
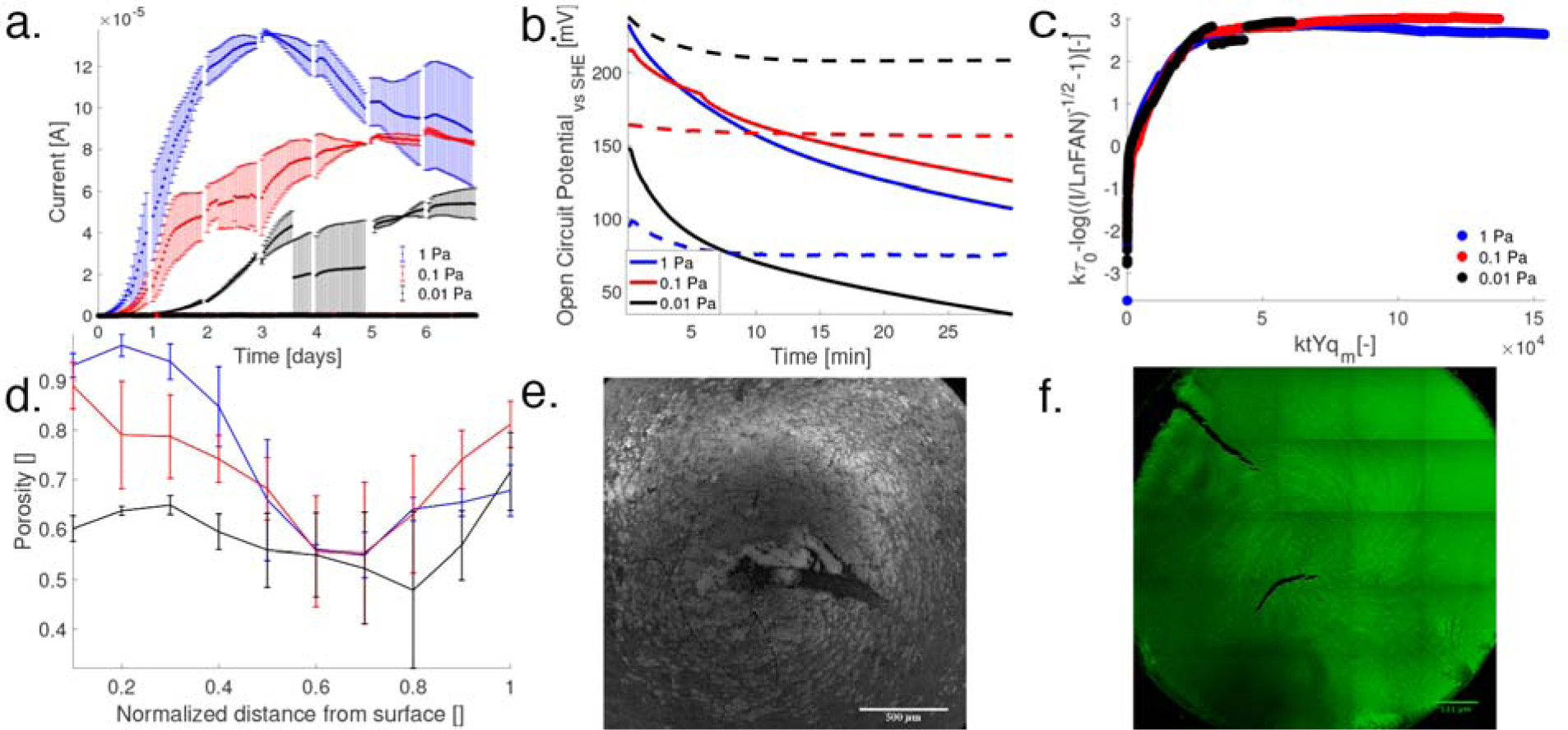
**a.** The current produced by *G. sulfurreducens* at three shear stresses of 1 Pa, 0.1 Pa (*s.e. n=3*) and 0.01 Pa (*s.e. n=2*). The abiotic measurements never exceed 10^−7^ A for each shear stress condition. The flux was fixed to an up-flow microbial fuel cell (He et al., ^*6*^) which also corresponded to the 1 Pa shear stress case. We find that at the lower shear stress, the maximum current persists longer. The maximum current was 136±1.24 μA, 89.3±1.24 μA and 54.03±7.08 μA at 1 Pa, 0.1 Pa and 0.01 Pa in blue, red and black respectively. **b.** Early time-averaged chronopotetiometry shows rapid decrease in current with bacteria present in contrast to ‘--’ lines of the abiotic control. **c.** Dimensional analysis reveals two parameters of interest, a scaled growth time, τ = *ktYq_m_*, and a ratio of current to maximum electrons delivered, 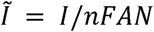, where *n* is the number of electrons transferred for a flux *N* to an electrode of area *A*, and *F* is Faraday’s constant. These parameters are fit to a logistic growth model, showing that the lowest shear case has not yet reached the maximum metabolic current output as would be predicted by scaling and would be anticipated from Figure 1a. This model does not account for the decrease in current. **d.** Porosity of the biofilm as a function of normalized distance from the surface obtained from confocal microscopy (Figure 1**f.**). This data shows that the biofilms in the 1 Pa and 0.1 Pa cases, which we predict as fully-developed, have similar structure, whereas the 0.01 lowest shear case does not. **e.** An image of the biofilm from an underperforming 1 Pa shear stress condition (16 μA after 7 days). The biofilm shows a growth pattern similar to the streamlines predicted by the von Kármán solution to flow at a rotating disk, SI Figure S4. **f.** A confocal image slice 6 μm from the surface of an electroactive biofilm taken after 7 days at 0.1 Pa shear stress, showing the predicted streamlines of the fluid flow which should not be present ~34 μm into the biofilm. We assume this is “memory” of flow-influenced adhesion.

We used dimensional analysis to estimate that the time to maximum current is decreased for increased shear rates from 9.8 days to 3.8 days, as shown in Figure 1c and Table 2. The electrochemical signal of the biofilm displayed similar characteristics to optical density measurements for planktonic cells^32^ and optical measurements of biofilm growth.^33^ Assuming that current is a proxy for metabolic rate,^2^ the initial lag phase, where little current is found, was seen in the 0.01 Pa case (Figure 1a). A rapid growth phase can be seen in all three shear rates tested, a stable growth phase can be seen in the 0.1 Pa case, and a decay phase can be seen in the 1.0 Pa case. Thus, we assumed that the dimensionless current should scale with dimensionless time according to a logistic growth model. Our results indicate a shear dependence on current not seen in the literature^19^ when both mass and shear were coupled (Figure 2a.) for the same species.

**Figure 2.**
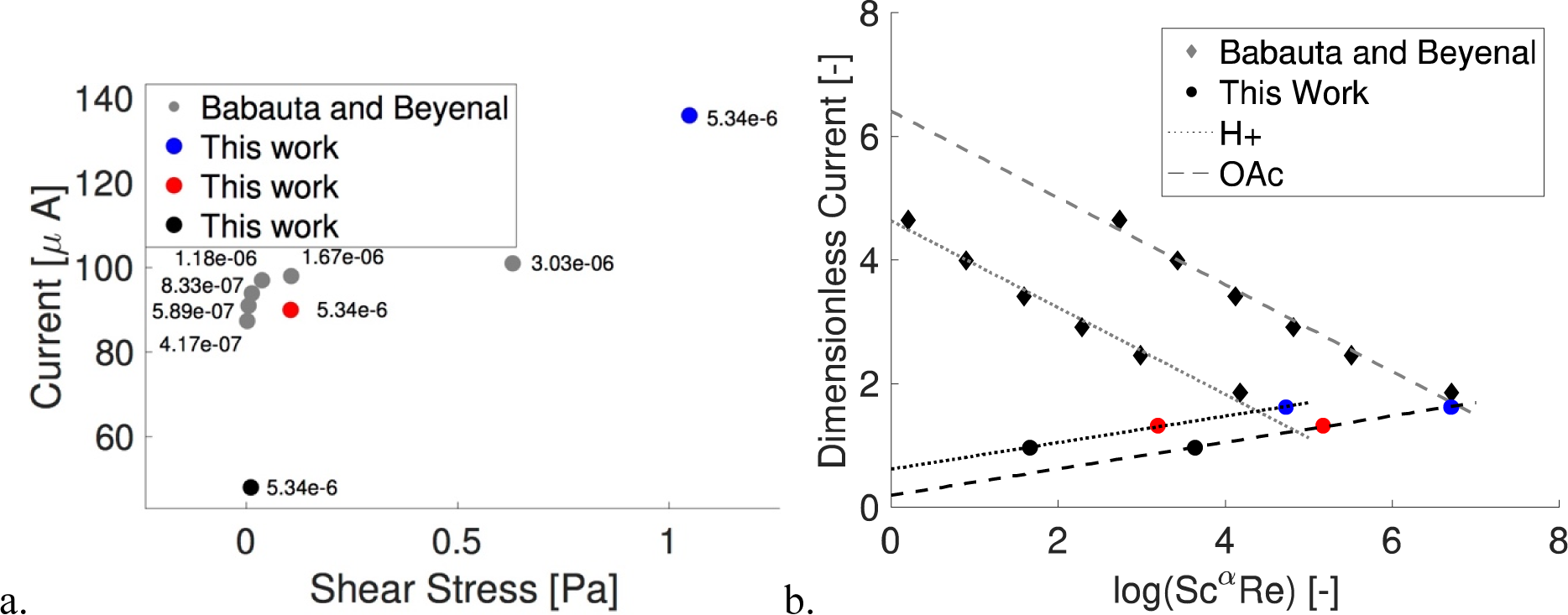
**a**. A comparison of shear stress and current found in our work with that found by Babauta and Beyenal using the same species, electron donor, and reactor dimensions. The flux in our work is strictly greater than their fluxes tested however the current results vary. 2.b. Dimensionless current against the product of the Schmidt number and Reynolds number shows linear dependence of biofilm metabolism on viscous shear and mass transport. The Schmidt number goes as a = 1 for both acetic acid and proton transport in this work, with *R^2^* = 0.99. The Scmidt number goes as α = −1 for acetic acid and α = 1.1 for proton transport for Babauta and Beyenal’s work, with *R^2^* = 0.98.

It is accepted that varying concentration (mass flux) changes metabolic (current) output for all bacteria, and varying electrode potential can improve power output in electroactive biofilms.^34^ From this hypothesis, previous studies have grown the biofilm under one shear stress condition and varied shear to see the impact on current.^19^ Using the product of Schmidt and Reynolds number as the common scaling for mass transport and the scaling of current from Figure 1 c., we find very different metabolic efficiencies and transport regimes for biofilms subjected to shear from different starting conditions. The maximum current is the same when scaled and occurs at the same transport condition whether looking at acetate (substrate) or protons (toxic waste byproduct), Figure 2 b for biofilms grown under shear or grown in static and then subjected to higher shear. However, biofilms grown in static conditions show a decrease in dimensionless current with increasing transport. Our dimensionless current is equivalent to the Coulombic efficiency, so we hypothesize that the biofilm is adapted to maximum utilization at its initial growth condition (shear stress of 0 Pa) and any increase is not beneficial. This adaptation is likely the higher porosity found for biofilms not grown under shear.^35^ For both, protons or acetate, biofilms in this work are strictly dependent on mass transport, i.e. the Peclet number. However, biofilm’s initially grown in static conditions are dependent on both diffusivity and viscosity for acetic acid transport and weakly dependent on viscosity for proton transport. While no hysteresis in current output was found for the biofilms grown in static conditions and then sheared indicating no loss of biomass,^19^ this dependence on viscosity should be further investigated. We caution that our dimensional model does not capture the decay shown in the high shear stress case, Figure 1 a., though it does help determine when it may be necessary to run experiments for longer durations.^1^

High metabolic rates described appear to come at the cost of sustained growth and stable metabolic activity. The highest shear stress, 1.0 Pa, produced the highest current density (see Table 2) yet immediately declined (Figure 1a). This decline in current was not present for the lower shear stress cases. While it has been shown that higher shear induces higher metabolic rates in non-electroactive biofilms,^36^ previous experiments were run for half the duration of the present study. As a result, they did not show the trade-offs found here-in for the highest shear.^36^ Measured electron diffusion current followed similar trends as the electrical current with shear. Since electron diffusion current is dependent on electron mediator concentration, bacteria concentration, and reaction rate,^37^ we hypothesize that the decline is due to either the death of bacteria or the loss of electron mediator activity. Existing hypotheses in the literature show that high current production^5^ and/or high metabolic activity^33^ leads to decreased pH in the biofilm,^5^ and that electron mediators/proteins lose activity at low pH. Electrochemical impedance spectroscopy, a measure of the complex resistance to charge transfer, similarly showed an increase in resistance upon a decrease in current (both electron diffusion and absolute), which could either imply cell death or loss of electron mediator activity, see SI Figure S2.

While biofilm health is negatively affected by higher shear stress over long-term growth, the biofilm retains memory of initial fluid flow conditions and does not change structure to counteract such stress, Table 3. The independence of thickness and shear for the two higher shear stress cases agrees with previous studies on membrane aerated biofilms.^10^ Since the lowest shear stress case has not reached maximal current, we conclude that biofilm thickness, surface roughness, and porosity are independent of shear for the conditions tested once the biofilm is fully developed (Figure 1d).

**Table 3.** Similarities in thickness (H) and surface roughness (*R_a_*) appear to confirm that the 1 and 0.1 Pa cases biofilms are fully developed while the 0.01 Pa case is still developing. *n = 2*

| Shear [Pa] | H [μm] | $R_a$ |
| --- | --- | --- |
| 1 | 41 | 2.50 |
| 0.1 | 41 | 2.07 |
| 0.01 | 16 | 1.36 |

The fluid streamline pattern (Figure 1f) is imprinted on the biofilm interior at a height of 6 μm, but not at the surface of the biofilm after seven days of growth, Figure S6. The stability of this interior structure is quantitatively seen in stable open circuit potential^31^ and redox potential,^37^ indicating a stable surface attachment of electrically respiring bacteria/proteins, SI Figures S3-4. The mechanisms of this imprinting were not studied, though it may be due to the transport of quorum sensing chemicals during biofilm formation. The memory of a complex substructure from initial growth conditions of a biofilm has been seen before,^11^ yet this is the first time to the authors’ knowledge that it has been shown under controlled initial conditions.

## Conclusion

While this study shows promising results for optimizing bioelectrochemical system startup times, it also shows biofilms’ limited ability to adapt to engineered environments. This result is consistent with the existing literature suggesting that shear stress does not induce changes in biofilm structure in both single and multi-species biofilms.^10,38^ Changes in structure and metabolism may offset each other when mass flux and shear stress are coupled, resulting in some of the conflicting reports on the independence of shear from both structure and metabolism.^19,39^ Individual bacteria cell size and transcriptomic changes in the biofilm should be measured in a homogenous biofilm over time to determine if there are other structural changes induced by shear not captured by macro-scale confocal imagery. The result that the biofilm has memory of the initial surface conditions may explain how nanotexture improves bacteria metabolism beyond increasing surface energy, by adding artificial porosity that can absorb metabolic byproducts.^40^ While modeling and experimentation has not produced physical laws that can predict biofilm porosity, there is increasing evidence that the formation of subsurface pores of the biofilm is governed by the initial surface forces. This is similar to how mammalian cell growth is governed by the surface tension of the substrate. We propose that dynamic control of shear stress should be considered an independent tool for biofilms to maintain reactor stability similar to variation of potential,^34^ batch feeding,^40^ and removal of “old” biomass.^29^

